# Accurate protein stability predictions from homology models

**DOI:** 10.1101/2022.07.12.499700

**Authors:** Audrone Valanciute, Lasse Nygaard, Henrike Zschach, Michael Maglegaard Jepsen, Kresten Lindorff-Larsen, Amelie Stein

**Affiliations:** Linderstrøm-Lang Centre for Protein Science, University of Copenhagen, Copenhagen, Denmark; Section for Computational and RNA Biology, Department of Biology, University of Copenhagen, Copenhagen, Denmark

## Abstract

Calculating changes in protein stability (ΔΔG) has been shown to be central for predicting the consequences of single amino acid substitutions in protein engineering as well as interpretation of genomic variants for disease risk. Structure-based calculations are considered most accurate, however the tools used to calculate ΔΔGs have been developed on experimentally resolved structures. Extending those calculations to homology models based on related proteins would greatly extend their applicability as large parts of e.g. the human proteome are not structurally resolved. In this study we aim to investigate the accuracy of ΔΔG values predicted on homology models compared to crystal structures. Specifically, we identified four proteins with a large number of experimentally tested ΔΔGs and templates for homology modeling across a broad range of sequence identities, and selected three methods for ΔΔG calculations to test. We find that ΔΔG-values predicted from homology models compare equally well to experimental ΔΔGs as those predicted on experimentally established crystal structures, as long as the sequence identity of the model template to the target protein is at least 40%. In particular, the Rosetta `cartesian_ddg` protocol is robust against the small perturbations in the structure which homology modeling introduces. In an independent assessment, we observe a similar trend when using ΔΔGs to categorize variants as low or wild-type-like abundance. Overall, our results show that stability calculations performed on homology models can substitute for those on crystal structures with acceptable accuracy as long as the model is built on a template with sequence identity of at least 40% to the target protein.

## Introduction

The vast majority of functions in a cell are carried out by proteins. These biomolecules typically need to fold into a stable tertiary structure in order to be functional. Experimental structure determination can provide us with high-resolution details of the arrangement of atoms in the folded protein, however it does not directly reveal the forces holding them together. These have historically been derived from mutagenesis experiments and measurements of their energetic consequences (Dill 1990; Gromiha et al. 1999; Nikam et al. 2021). From these studies we know that many changes are detrimental for protein stability (Tokuriki and Tawfik 2009), although stabilizing mutations have also been observed (Shoichet et al. 1995). In the absence of experimental data, the stability of variant proteins is commonly assessed by computational means with so-called assessment of the change in thermodynamic stability upon mutation (ΔΔG). This assessment has many practical applications, from protein engineering (Goldenzweig and Fleishman 2018) to identification of disease-causing mutations (Yue, Li, and Moult 2005; Stein et al. 2019). Hence, a number of methods for ΔΔG prediction have been developed. Among the most widely used ones are FoldX (Guerois, Nielsen, and Serrano 2002; Radusky and Serrano 2021; Gerasimavicius, Liu, and Marsh 2020) and Rosetta (Kellogg, Leaver-Fay, and Baker 2011; Park et al. 2016; Frenz et al. 2020).

These methods require a structure of the wild-type protein as input, which in combination with molecular force fields or energy functions in principle allows for the generation of accurate and high-resolution models of the mutant protein. At the same time, though, this requirement represents an important limiting factor for their application. Recent advances in sequencing technology particularly highlight the gap between known sequences and known structures; many new protein families have been found in large-scale sequencing experiments for which no structural information is available, and there are vast differences in coverage between species (Marks, Hopf, and Sander 2012). For proteins where the structure of another family member is known, homology modeling can be employed to provide a computationally-derived structure model. Briefly, this approach relies on using the solved structure of a homologue of the protein of interest as a template for the backbone of the target sequence, and using loop modeling and refinement of intramolecular interactions by either physics-based or empirical force fields to generate a high-resolution model (Martí-Renom et al. 2000). For template-target pairs with sequence identity >70%, models are typically fairly accurate (1Å–2Å RMSD), whereas pairs below 30% sequence identity are in the so-called “twilight zone” and often have different 3D structures and distinct functions (Martí-Renom et al. 2000). However, calculations of the change in thermodynamic stability upon mutation require accurate backbone as well as side chain placement, and the latter may require a higher identity between template and target of around 50% (Chung and Subbiah 1996). Model quality depends on the template-target sequence identity. A number of tools have been developed to assess the quality of an individual homology model based on similarity to experimentally resolved structures in terms of torsion angles, core packing and other high-resolution properties (Kryshtafovych et al. 2016; Shen and Sali 2006)

If ΔΔG calculations could be performed reliably on computationally generated models of protein structure, this would greatly extend the scope of their applicability, and open up e.g. possibilities of structure-based re-engineering of proteins without an experimentally determined structure. Only about 15% of the human proteome is covered by experimental structures, but with inclusion of homology models, coverage could be doubled or even quadrupled, depending on whether templates with down to 50% or 20% sequence identity are chosen, respectively (Bienert et al. 2017). Both FoldX and Rosetta, however, were developed and benchmarked for use with experimentally determined structures so the question remains how applicable they are to computationally-derived structures.

On one hand, stability effects have been shown to be largely consistent when introducing a particular mutation into different homologs in the same family (Ashenberg, Gong, and Bloom 2013). On the other hand, ΔΔG predictions have been shown to suffer from challenges in reversibility, i.e., the predicted ΔΔG of a V→A mutation is not the inverse of the corresponding A→V mutation, revealing gaps in capturing the underlying biophysical forces and a substantial dependency on the structural input model (Thiltgen and Goldstein 2012; Johansson et al. 2016).

Here we set out to assess whether ΔΔGs can be accurately predicted using homology models, and to what extent the accuracy is dependent on the sequence identity between template and target. To assess the dependence on target-template similarity systematically, we chose four proteins for which a substantial number of biophysical ΔΔG measurements as well as template protein structures across a range of sequence identities are available. From those four proteins we established a dataset of 344 mutations. For an independent comparison, we also calculate full-saturation ΔΔGs for ~3,500 single amino acid variants of a protein assayed for cellular abundance (Matreyek et al. 2018), again using experimental structures, close and distant homology models as starting points. Overall, we find that all tools work well on experimentally resolved structures, and that Rosetta `cartesian_ddg` performance appears robust on homology models with at least 50% sequence identity.

## Results and Discussion

Our goal was to determine the accuracy of predicted stability changes in proteins calculated using homology models instead of experimentally determined structures. To this end, we compare a set of experimentally-determined stability changes (ΔΔG) to predicted ΔΔG values calculated on either experimentally determined structures or homology models

For the experimental data we used a recently curated database of 2971 experimentally-determined stability changes (ΔΔG) (Ó Conchúir et al. 2015) which is largely based on the Protherm database (Gromiha et al. 1999). From this set we selected four different proteins based on the following criteria: First, each protein should have a large number of experimentally determined single point ΔΔG values. Second, a high-resolution crystal structure of the wild-type protein should be available. Finally, we required that for each protein a number of homologues with varying degrees of sequence identity existed, which themselves had experimentally-resolved structures. This latter criterion is central to our objective as it enables us to build homology models of the four proteins using templates with varying degrees of sequence identity. With these criteria we selected human lysozyme (PDB ID 1LZ1), the barley chymotrypsin inhibitor CI2 (2CI2), *E. coli* RNAseH (2RN2) and sperm whale myoglobin (1BVC) which together have 344 experimentally-determined ΔΔG-values.

For the computationally-derived data, homology models with increasing divergence from the wild-type protein were needed. We divided the range from 100% to 25% sequence identity into 15% intervals and obtained the following five bins: bin1: (100%,85%]; bin2: (85%,70%]; bin3: (70%,55%]; bin4: (55%,40%]; bin5: (40%,25%]. For each protein and each bin we searched the PDB for homologues and picked the one with a sequence identity closest to the middle of the bin. For example, in the case of sperm whale myoglobin, bin2 was populated using an elephant myoglobin (PDB ID: 1EMY) with a sequence identity of 81% to the sperm whale protein. In this way we populated the full matrix of structures/sequences covering all four proteins and all five bins, apart from bin3 for CI2 for which we could not find a suitable template (Suppl. Table S1).

We used Modeller (Webb and Sali 2016) to build homology models for all four proteins using the 19 different templates. Modeller is widely used and, importantly, allows for direct selection of the template, which ensures that the resulting model’s quality was not influenced by the availability of a crystal structure of the wild-type protein. We show the modeling results for sperm whale myoglobin in Fig. 1 (top row) where we overlay the models to the structure of the wild-type protein; corresponding plots for the remaining proteins can be found in the supporting online material (Suppl. Fig. S1-S3). As expected, the agreement between the crystal structure of the wild-type protein and the template-based models decreases as the sequence of the template diverges from that of the wild-type protein.

**Fig. 1:**
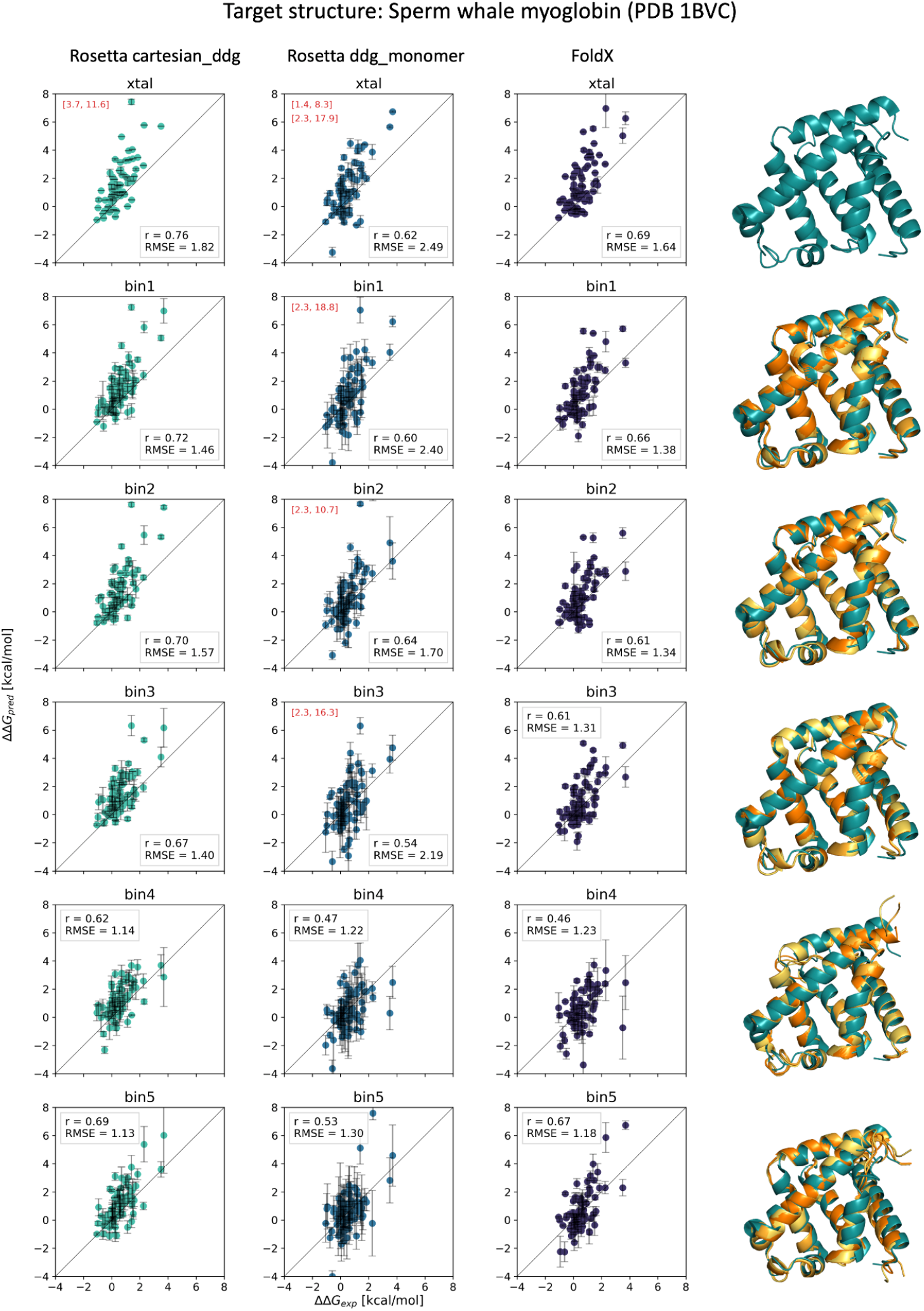
Comparison of computationally-derived ΔΔG values with experimental values on sperm whale myoglobin using three different ΔΔG-prediction methods and either the crystal structure or a homology model. Each field shows a scatter plot comparing predicted and experimental ΔΔG values, including the corresponding Pearson’s correlation coefficient (r) and the root mean square error (RMSE; kcal/mol). If present, values outside the [-4,8] kcal/mol range on either axis are listed in red in the top left corner of each plot. First row: ΔΔG values computed on the crystal (xtal, 1BVC) structure of the wild-type protein. Following rows: ΔΔG values computed on homology models with decreasing sequence identity between target and template. First column (turquoise): ΔΔG values predicted by Rosetta `cartesian_ddg`. Second column (blue): ΔΔG values predicted by Rosetta `ddg_monomer`. Third column (violet): ΔΔG values predicted by FoldX. Last column: Superimposition of the wild-type protein crystal structure with the homology model. Agreement between the crystal structure and the model deteriorates as the sequence identity between template and target decreases (top to bottom, bin1 = highest identity).

We then calculated ΔΔG values with three different structure-based methods: FoldX, Rosetta `ddg_monomer` and Rosetta `cartesian_ddg`. All three methods are widely used for ΔΔG-predictions but, to our knowledge, have never been benchmarked for use with template-based models. For each method and protein we ran ΔΔG calculations on each sequence identity bin plus the wild-type crystal structures and thus predicted the 344 ΔΔG-values corresponding to the protein variants we have experimental data for.

We compared predicted and experimental ΔΔG-values using the Pearson correlation coefficient (r) and the mean absolute error (MAE). For MAEs to be comparable across methods, we first calculated a scaling factor to bring the predicted ΔΔG values to kcal/mol (see Methods). The results are illustrated in Fig. 1 with sperm whale myoglobin as an example. First, we find that all three methods give a comparable agreement to the experimental data when we use a high-resolution crystal structure of the wild-type protein as input for stability predictions (row 1). As expected, we also see that the predicted ΔΔG-values correlate less with experimental data as the sequence identity of the template used to build the homology model decreases (following columns). For example, when we use FoldX on either the crystal structure of sperm whale myoglobin (wild-type structure) or the homology model build on wild boar myoglobin (bin1; 86% sequence identity), we obtain r ~ 0.7 between the predicted and experimental ΔΔG-values. This drops to r ~ 0.6 for bin2 (Asian elephant myoglobin; 81%) and bin3 (Sea turtle myoglobin; 65%) and further for bins 4 and 5. A similar trend was observed for all three methods and all four proteins (Figs. 1 and S1–S3), though with apparent differences in robustness towards using homology models.

As a baseline for minimal expected ΔΔG prediction performance, we developed two simple null models, which are designed to be independent of protein structure and the specific sequence. Both null models are based on the curated experimental ΔΔG database mentioned above (Ó Conchúir et al. 2015). The first null model simply assigns the average over all of these experimental measurements, ΔΔG=1.15 kcal/mol, to *any* mutation. Since most stability changes in the set of 344 mutations that we examine are of this order of magnitude, this model achieves a MAE=1.1 kcal/mol to the experimental ΔΔG-values. However, the Pearson correlation is very poor (r=0) since this model assigns the same value to all mutations. To introduce more specificity, we designed a second null model in which we again use the average ΔΔG over the entire database, but split up by wild-type and mutant residue, so that we obtain 380 averages corresponding to the 380 possible pairs of the 20 different wild-type amino acids with their 19 possible substitutions (Fig. 2). Before applying this null model to our set of mutations, we in practice make two modifications. First, since the 344 ΔΔG-values we use for validation are a subset of the 2971 values in the full dataset and went into the averages, we leave out all the ΔΔG-values pertaining to a protein when we apply that null model to that protein. This is particularly important for the many fields in the null model’s 20×19 matrix that include only very few mutations. Second, we use the global average of ΔΔG=1.15 kcal/mol for any mutations that are not present in the original dataset after leaving out the mutations in the protein that we apply the model to, such as histidine to methionine. When we use this modified null model to predict the stability outcome of the 344 mutations we achieve MAE=1.1 kcal/mol and r=0.42.

**Figure 2:**
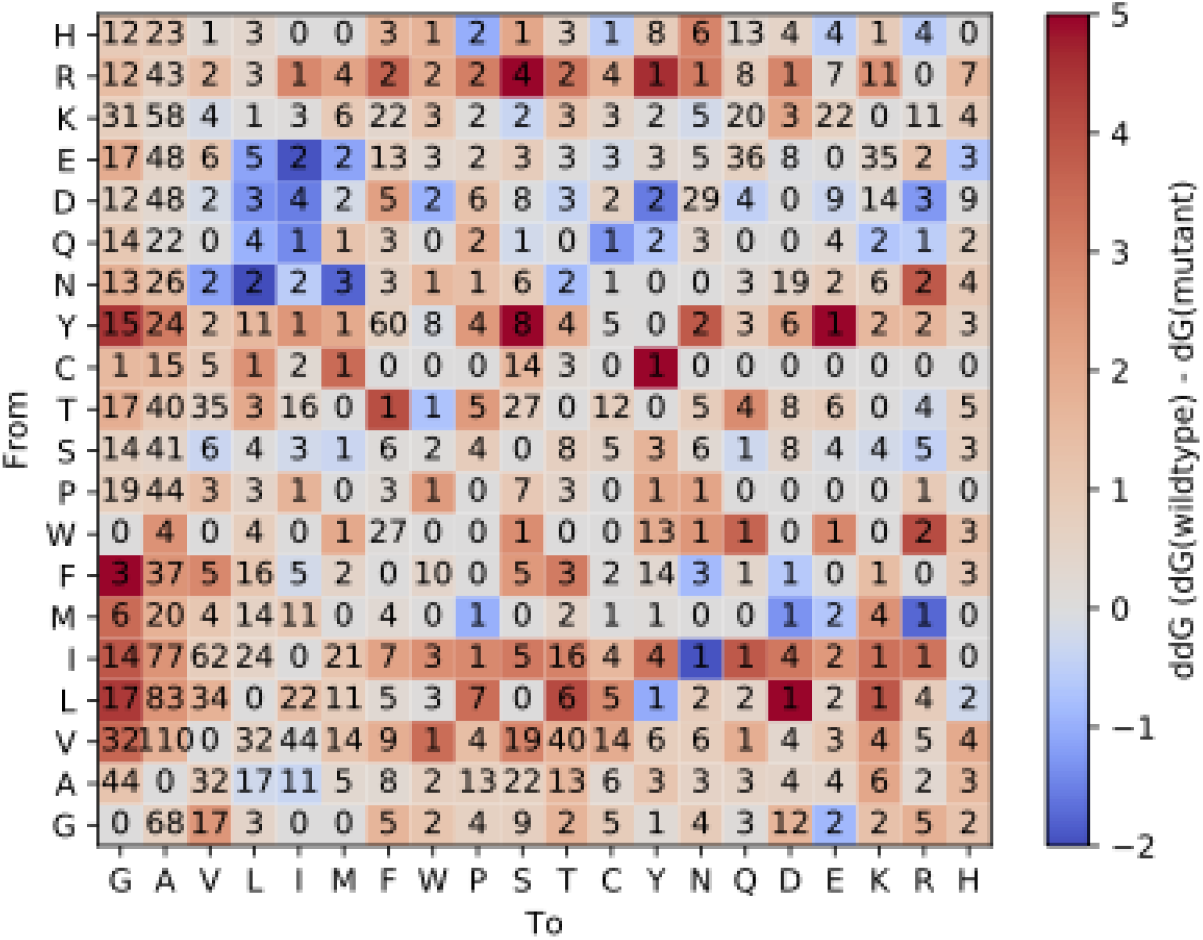
Mutations in the curated ProTherm dataset split by wild-type and target amino acid. These data are used in the null models and are a superset of the test data. Each entry lists the number of mutations of that type, and the color indicates the average change in stability (in kcal/mol).

With the (second) null model in hand, we then compared the results from all three structure-based methods and aggregated the results over all four proteins (Fig. 3). As previously observed for myoglobin (Fig. 1), we find that all three methods perform comparably when using the crystal structure of the wild-type proteins as input. Looking at the correlation coefficients, however, we also find that the methods appear to display different levels of sensitivity to the input structure. Both FoldX and Rosetta `ddg_monomer` appear more sensitive to the input structure than `cartesian_ddg` since the correlation of ΔΔG-values from homology models with the experimental ones is substantially lower than when using the wild-type crystal structures. In particular, when using the homology models from bins 4 and 5 (which in practice corresponds to sequence identities <45%) both FoldX and Rosetta `ddg_monomer` give results that are at most only slightly better than the (second) structure-independent null model. This decrease in accuracy is also reflected in an increase in the MAE, though we focus less on this value as it is less sensitive to the variation in structure and is also well-captured by the null models. In contrast to that, the results from Rosetta `cartesian_ddg` appear substantially more robust against changes in the input structure. Both the correlation coefficient and MAE vary little across bins 1 to 4 and are comparable to those obtained when using the wild-type crystal structures as input. To summarize, in practice we find that at least for the mutation dataset we used, Rosetta `cartesian_ddg` predicts stability changes with similar accuracy on homology models as when using crystal structures of the actual proteins so long as the template sequence identity is > 40%.

**Figure 3:**
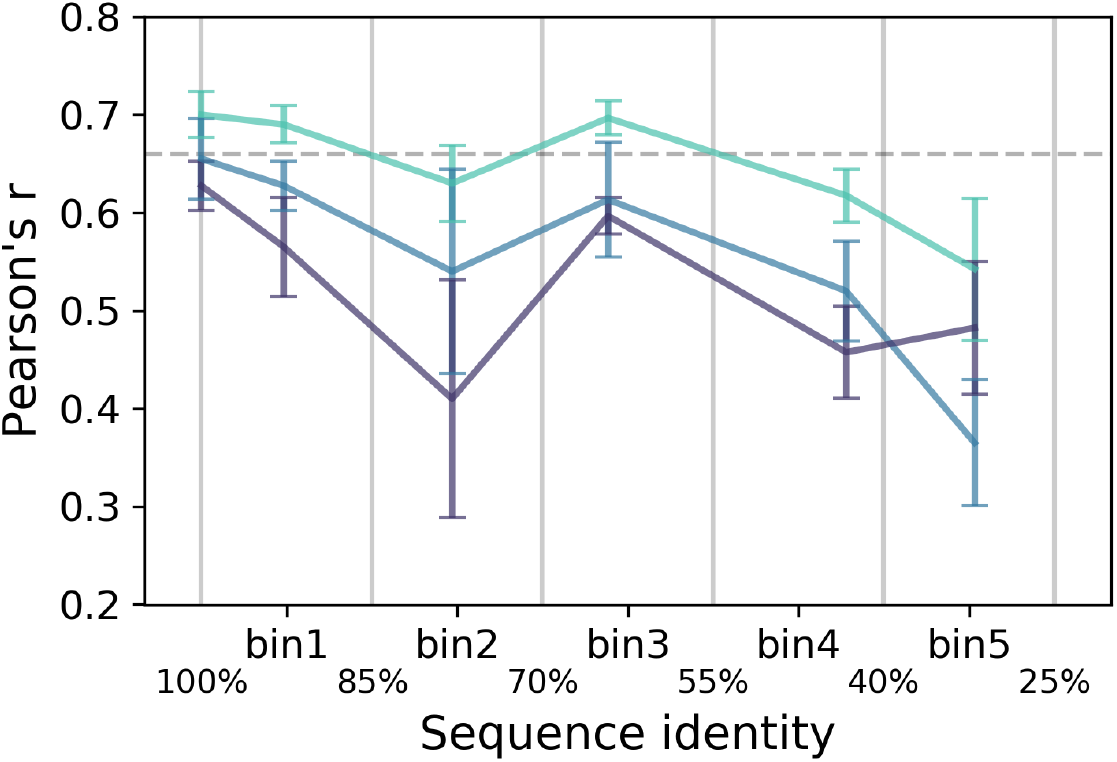
Correlation of ddGs calculated on the original crystal structure as well as homology models with decreasing sequence identity to the experimentally determined values. Turquoise, Rosetta `cartesian_ddg`, blue, Rosetta `ddg_monomer`, violet, FoldX. The dashed line indicates the average performance on the original crystal structures. Note that bin2 contains two models based on NMR structures, see also Suppl. Fig. S4.

The results detailed above depend on the availability of the crystal structures and homology models of the four proteins, as well as the experimental ΔΔG measurements. As an alternative approach, and to check for robustness, we also calculated correlation coefficients using the ΔΔG values calculated from the crystal structures as the reference instead of the experimental ones (Fig. S4). Per definition, the crystal structures have r=1, and hence the correlation coefficients from this analysis are higher than those using the experimental data as reference. This observation is expected given remaining biases in stability prediction methods. Nevertheless, the results we obtain here confirm the previous results, namely (i) a general decrease in the accuracy of stability predictions as homology models are based on increasingly distant templates and (ii) Rosetta `cartesian_ddg` appearing less sensitive than the other two methods.

The key result obtained above is that the Rosetta `cartesian_ddg` approach to predict stability changes appears to be relatively insensitive to minor differences in the input structure so that it gives fairly accurate results also when applied to template-based models from Modeller. The analyses are, however, based on a biased set of mutations with many mutations being hydrophobic deletion mutations and a high proportion of mutations to alanine or glycine (similar to Fig. 2). For a more unbiased set of single substitutions, we turned to recent data from a type of multiplexed assay of variant effects (MAVE) experiment termed VAMP-seq, performed on a protein called TPMT(Matreyek et al. 2018). Briefly explained, a MAVE involves creating a library of variants and selecting for a property of interest such as cell growth or fluorescence of a reporter protein (Fowler, Stephany, and Fields 2014). In a VAMP-seq experiment the library of protein variants are fused to green fluorescent protein (GFP) and expressed in cell culture. The brightness of each cell’s GFP fluorescence will be a function of the abundance of the protein variant (Matreyek et al. 2018). By sorting cells based on their fluorescence, and using DNA sequencing to determine the frequency of every variant before and after sorting, a single experiment provides abundance data for thousands of variants. Since destabilized proteins are degraded by protein quality control, a variant protein that has high abundance is expected to have little changes or only stabilizing changes to its thermodynamic stability (ΔΔG). Vice versa, a variant protein with low cellular abundance may have large changes in its thermodynamic stability compared to the wild-type protein (though proteins can also be degraded for other reasons). We use this relation to evaluate whether ΔΔG values calculated on either the crystal structure or the homology models behave in accordance with the experimental abundance data. In principle the VAMP-seq results are semi-quantitative, but we only use the information whether variants were low- or high-abundance.

In addition to the crystal structure of wild-type TPMT (PDB ID 2BZG) we built homology models of TPMT using templates with 80% identity (2GB4) and 32% identity (3LCC and 1PJZ). Since the VAMP-seq experiment does not directly probe thermodynamic stability, we instead used a ‘receiver operating characteristic’ (ROC) curve analysis to relate ΔΔG values and abundance classification. Specifically, we vary a cut-off for the predicted ΔΔG values, and assess how well it divides the variant proteins into low and high-abundance in terms of true positive rate (sensitivity) and false positive rate (specificity). The ‘area under the curve’ (AUC) statistic assesses how well the predicted ΔΔG values balance sensitivity and specificity across different thresholds. A ROC curve along the diagonal corresponds to a random prediction method and has AUC=0.5, whereas a method that can perfectly separate low- and high-abundance variants would have AUC=1. Our results show that ΔΔG values from Rosetta `cartesian_ddg` achieve a rather high accuracy both when using the wild-type TPMT structure (AUC=0.87) and the model based on 2GB4 (AUC=0.88) (Fig. 4). Also, as expected we find decreased prediction accuracies when using the two models based on templates with 32% sequence identity (AUC=0.78 and 0.77, for 3lcc and 1pjz respectively). Thus, the results obtained using these VAMP-seq data confirm the previous observation from the comparison to experimental ΔΔG values, namely that Rosetta `cartesian_ddg` provides relatively accurate predictions, and that there is little deterioration when the input structure is based on a close homologue. As the VAMP-seq library was designed to target every possible single amino acid substitution, it is much more balanced than data from ProTherm, which is dominated by alanine scanning results (Stein et al. 2019).

**Figure 4:**
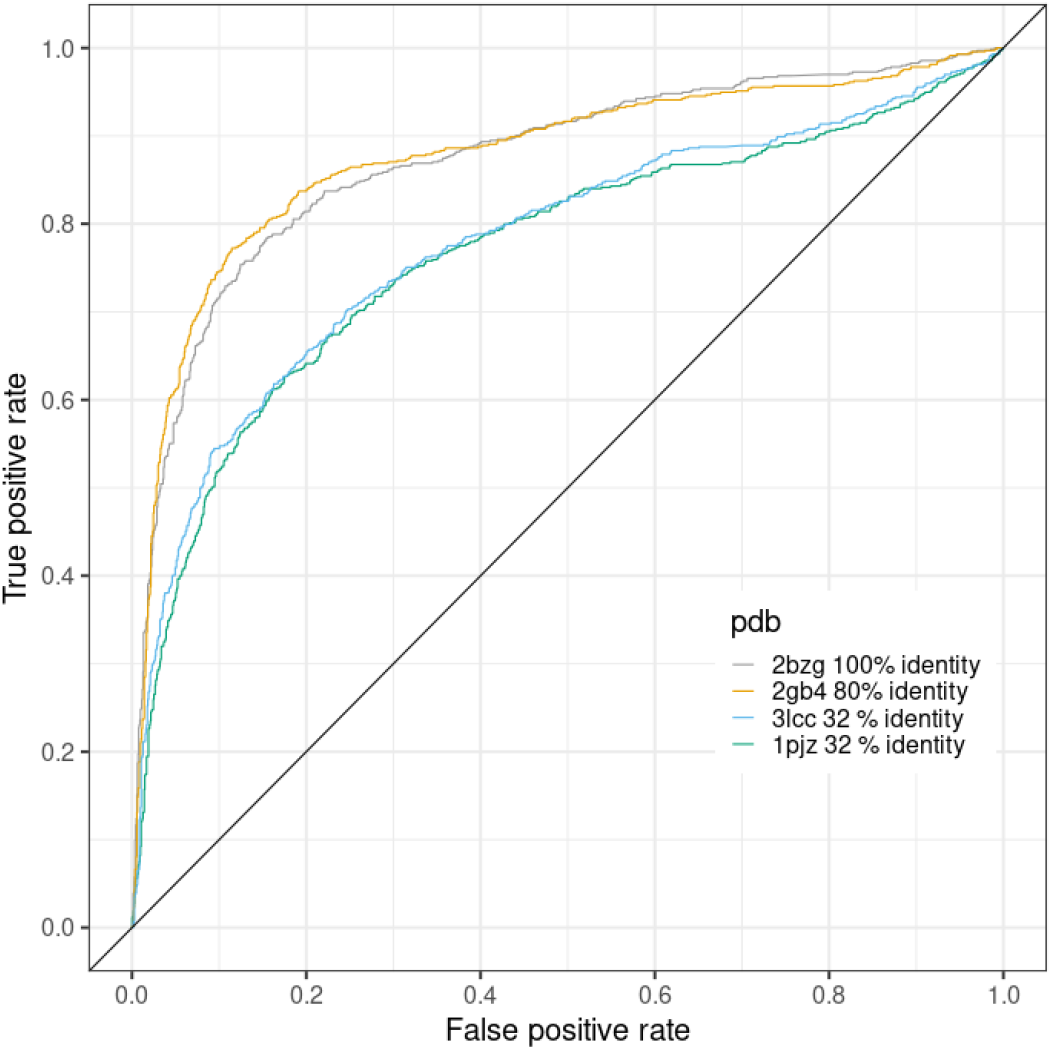
Receiver Operating Characteristic (ROC) curve of classifying variants of human thiopurine methyltransferase (TPMT) into high and low abundance based on the ΔΔG calculated using Rosetta `cartesian_ddg`, compared to experimentally determined abundance by VAMPseq ((Matreyek et al. 2018)). ΔΔG values were calculated on a crystal structure of TPMT (2BZG, gray, AUC=0.873), a homology model built on a template with 80% sequence identity (2GB4, orange, AUC=0.877), and two homology models built on templates with 32% sequence identity each (3LCC, blue, AUC=0.877; 1PJZ, cyan, AUC=0.768). Performance of a random model is indicated by the diagonal line (AUC = 0.5).

## Conclusions

The ability to predict changes in thermodynamic stability upon single amino acid substitutions in proteins is important in a wide range of practical and scientific problems. FoldX and the two Rosetta applications `ddg_monomer` and `cartesian_ddg` are widely used for that. These methods were parameterized and benchmarked using high-resolution crystal structures, and are recommended to be used with such structures as input. Many proteins of interest however do not have crystal structures available and here template-based (homology) modeling provides an appealing alternative. We asked how accurate the three methods are when used on homology models, and how the results depend on the sequence identity of the template used in model building. While the template-dependency of homology modeling is well-studied, it has to our knowledge not been investigated how sensitive computational stability prediction methods are to the structural noise induced by such models.

We therefore devised and applied a strategy to test how well these three methods work outside their normally recommended application range. Specifically, we chose four proteins for which we have extensive experimental stability measurements, crystal structures of the wild-type protein and templates of varying sequence identities to build homology models. Our analyses show that all three methods perform comparably well when applied to the crystal structures of the four proteins (Fig. 3). Further, the Rosetta `cartesian_ddg` method appears to be relatively robust to structural noise and errors introduced by homology modeling, at least for models based on templates with sequence identities greater than 40% to the target protein (Fig. 3). This observation was corroborated by using a ROC analysis to relate predicted stability measurements to high-throughput abundance measurements from VAMP-seq on the protein TPMT, though in this case we could only apply the analysis to a smaller number of templates (Fig. 4).

We further devised two simple null models for stability predictions that effectively require a look-up in a table of average experimental values. While we do not recommend using this model to perform predictions in practical applications, we suggest that it is a useful baseline for ΔΔG benchmark evaluation in general.

Overall, our results suggest that it should be possible to extend the range of applicability of structure-based protein stability predictions to homology models and thus to a much larger number of sequences. For example, about 15% of the human proteome is covered by experimental structures, however the SWISS-MODEL repository contains structural information for ~30% of the human proteome if one includes homology models with template sequence identities > 40% (Bienert et al. 2017).

We conclude by suggesting that these kinds of analyses should be applied more broadly, and to a wider range of data and proteins, and point out a recent assessment of performance of multiple methods on AlphaFold2 (Jumper et al. 2021) models, compared to measurements from MAVE experiments (Akdel et al. 2021). Also, it is our hope that future developments of stability prediction methods, as well as other methods to analyze protein structures, will be developed with applications on homology models or predicted structures in mind. Thus, we suggest it should be possible to develop methods that are even more robust to structural noise and possibly supported by structure-independent methods. If we can increase robustness down to templates with about 30% sequence identity we should be able to analyze about half of the human proteome with these methods.

## Methods

### Homology Modeling

Homology models for human lysozyme, chymotrypsin inhibitor CI2, *E. coli* RNAseH and sperm whale myoglobin were generated using Modeller’s automodel class (Webb and Sali 2016). The target sequences for the four proteins were obtained from Uniprot (The UniProt Consortium and The UniProt Consortium 2017). Templates were obtained from a search of the pdball database (obtained from https://salilab.org/modeller/supplemental.html). The search results were binned according to sequence identity in 15% intervals and we selected the template closest to the middle of the interval for each respective bin. For the model building, all settings and parameters were left at their default value. 20 models were generated for high sequence identity bins (>55%) and 1000 models for low sequence identity bins (<55%). Three models were selected by the lowest DOPE scores in each sequence identity bin.

### ΔΔG calculations: FoldX

FoldX version 3.0b4 was used. Prior to ddg prediction, the structures were run through the RepairPDB function, as recommended in the FoldX documentation. The FoldX ddg predictions were performed using the BuildModel command of FoldX (Guerois, Nielsen, and Serrano 2002). For crystal structures, 5 iterations were used as recommended. For homology models, 2 BuildModel iterations were carried out on each of the top 3 models, and the average across those 6 ΔΔGs is reported.

### ΔΔG calculations: Rosetta ddg_monomer

We used Rosetta from October 2018 with git SHA1 `ce9cb339991a7e8ca1bc44efb2b2d8b0a3d557f8`. We ran `ddg_monomer` with the global repacking option (protocol 16, (Kellogg, Leaver-Fay, and Baker 2011)) and the talaris2013 energy function, which has been shown to improve ΔΔG calculation performance (O’Meara et al. 2015; Ó Conchúir et al. 2015). We carried out initial constrained minimization as recommended (Kellogg, Leaver-Fay, and Baker 2011). Average ddGs of 50 iterations are reported for the crystal structure. For homology models, the top 3 models were chosen, 20 ΔΔG iterations were performed on each, and the overall average is reported.

Flags for constrained minimization:

~~~
-in::file::fullatom
-ignore_unrecognized_res
-fa_max_dis 9.0
-ddg::harmonic_ca_tether 0.5
-ddg::constraint_weight 1.0
-ddg::sc_min_only false
~~~

Flags for `ddg_monomer`:

~~~
-ddg::weight_file soft_rep_design
-ddg::local_opt_only false
-constraints::cst_file ca_dist_restraints.cst
-fa_max_dis 9.0
-ddg::min_cst true
-ddg::iterations 20
-ddg::mean false
-ddg::min true
-ddg::sc_min_only false
-ddg::output_silent true
-ddg::ramp_repulsive true
-ddg::mut_only
~~~

### ΔΔG calculations: Rosetta cartesian_ddg

We used Rosetta from October 2018 with git SHA1 `ce9cb339991a7e8ca1bc44efb2b2d8b0a3d557f8`. For `cartesian_ddg` (Park et al. 2016; Frenz et al. 2020) we first performed constrained relaxation, then carried out 3 iterations of the `cartesian_ddg` protocol, and reported average ΔΔGs over these. For homology models, we carried out one iteration each over the 3 top homology models, and reported the average over these.

Flags for relax:

~~~
-relax::constrain_relax_to_start_coords
-ignore_unrecognized_res
-missing_density_to_jump
-ex1 -ex2
-relax::min_type lbfgs_armijo_nonmonotone
-flip_HNQ -no_optH false
-relax::coord_constrain_sidechains -relax::cartesian
-beta
-score::weights beta_nov16_cart
~~~

Flags for cartesian_ddg:

~~~
-fa_max_dis 9.0
-ddg::dump_pdbs false
-ddg::iterations 1
-score::weights beta_nov16_cart
-missing_density_to_jump
-ddg::mut_only
-ddg::bbnbrs 1
-beta_cart
-ex1 -ex2
-ddg::legacy false
-optimize_proline
~~~

### Scaling to kcal/mol

For each method a scaling factor was established to describe the relationship between the energy units reported by the method and kcal/mol. This was done by comparing the values predicted on the high quality crystal structures of the wild-type proteins against the experimental values and then forcing a linear fit through 0. The slope of this fit is the method’s scaling factor which was applied to all predictions made by the method. We obtained the following scaling factors:

FoldX: 1/1.039

Rosetta `cartesian_ddg`: 1/2.469 - note that a factor of 1/2.9 has previously been reported (Park et al. 2016)

Rosetta `ddg_monomer`: 1/1.233

### Null models

Data was extracted from a curated version of ProTherm (Ó Conchúir et al. 2015). In addition to the published curation, we noticed two apparent self-mutations with non-zero ΔΔG, T53T in protein G and K33K in ubiquitin. We traced both back to articles reporting the ΔΔG with reference to the stability of the sequence with alanine at the position of interest (Smith, Withka, and Regan 1994; Loladze and Makhatadze 2005), as opposed to the more common convention of reporting with respect to wild-type. We removed entries from both articles from the dataset before creating the null models.

For the null models we used the average experimental ΔΔG value of the given amino acid substitution across all the proteins in ProTherm, excluding the values from the protein itself.

### ROC curves

ROC curves were drawn and AUCs calculated with the Rpackage ROCR, directly using ΔΔG values obtained from running Rosetta `cartesian_ddg` on the homology models as predictor variables and low and high abundances classes from (Matreyek et al. 2018) as the two underlying true classes.

## Acknowledgements

We thank members of the Lindorff-Larsen and Stein labs for helpful discussions and comments. This work is supported by a Novo Nordisk Foundation Challenge Grant (PRISM, NNF18OC0033950 toK.L.-L., A.S.) and the Lundbeck Foundation (R272-2017-4528 to A.S.).

## Supplemental Material

**Supplemental Table S1:** Structures used for ΔΔG calculations and as templates in homology modeling.

| Target protein |  | bin1 | bin2 | bin3 | bin4 | bin5 |
| --- | --- | --- | --- | --- | --- | --- |
| Human lysozyme 1LZ1/A | Template PDB ID | 2BQB/A | 1IVM/A | 5V92/A | 4YF2/A | 1HFX/A |
|  | Resolution | 1.8 Å | NMR | 1.11 Å | 2.15 Å | 1.9 Å |
|  | Coverage | 100% | 100% | 99% | 98% | 95% |
|  | Seq identity | 98% | 78% | 62% | 47% | 33% |
| Barley chymotrypsin inhibitor CI2 2CI2/I | Template PDB ID | 1TM5/I | 1CIS/A |  | 3RDY/A | 1SIB/I |
|  | Resolution | 1.45 Å | NMR | - | 1.84 Å | 2.4 Å |
|  | Coverage | 97% | 98% |  | 100% | 95% |
|  | Seq identity | 91% | 82% |  | 40% | 34% |
| *E. coli* RNase H 2RN2/A | Template PDB ID | 2Z1J/A | 1JL2/A | 2ZQB/A | 3H08/A | 3HST/B |
|  | Resolution | 2.38 Å | 1.76 Å | 2.5 Å | 1.6 Å | 2.25 Å |
|  | Coverage | 97% | 96% | 95% | 89% | 81% |
|  | Seq identity | 90% | 72% | 67% | 43% | 30% |
| Sperm whale myoglobin 1BVC/A | Template PDB ID | 5YCI/A | 1EMY/A | 1LHS/A | 1MYT/A | 1V5H/A |
|  | Resolution | 1.97 Å | 1.78 Å | 2 Å | 1.74 Å | 2.4 Å |
|  | Coverage | 100% | 100% | 100% | 94% | 95% |
|  | Seq identity | 92% | 81% | 65% | 43% | 31% |

**Supplemental Figure S1:**
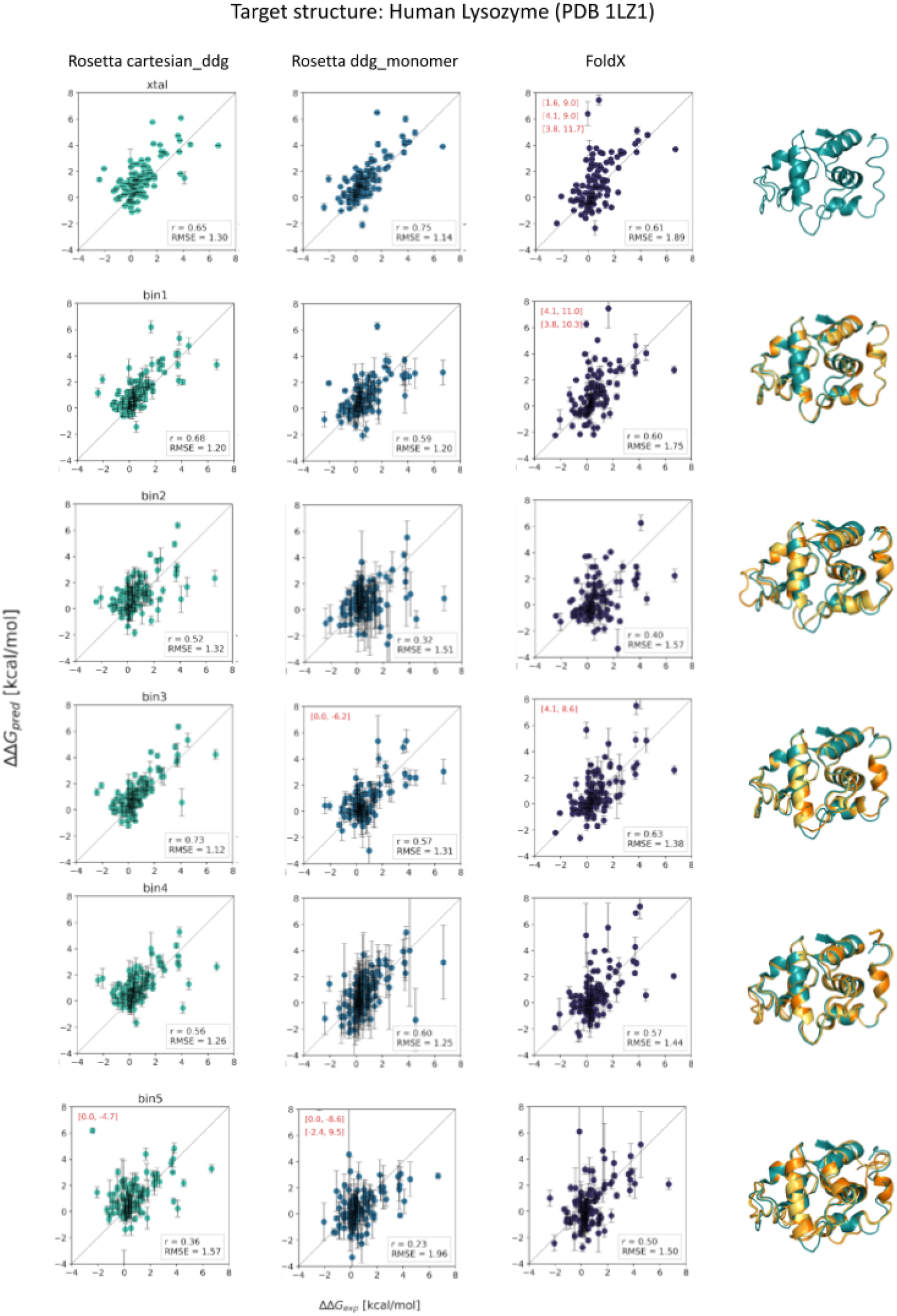
Comparison of computationally-derived ΔΔG values with experimental values on human lysozyme using three different ΔΔG-prediction methods and either the crystal structure or a homology model. Analogous to Figure 1 in the main manuscript.

**Supplemental Figure S2:**
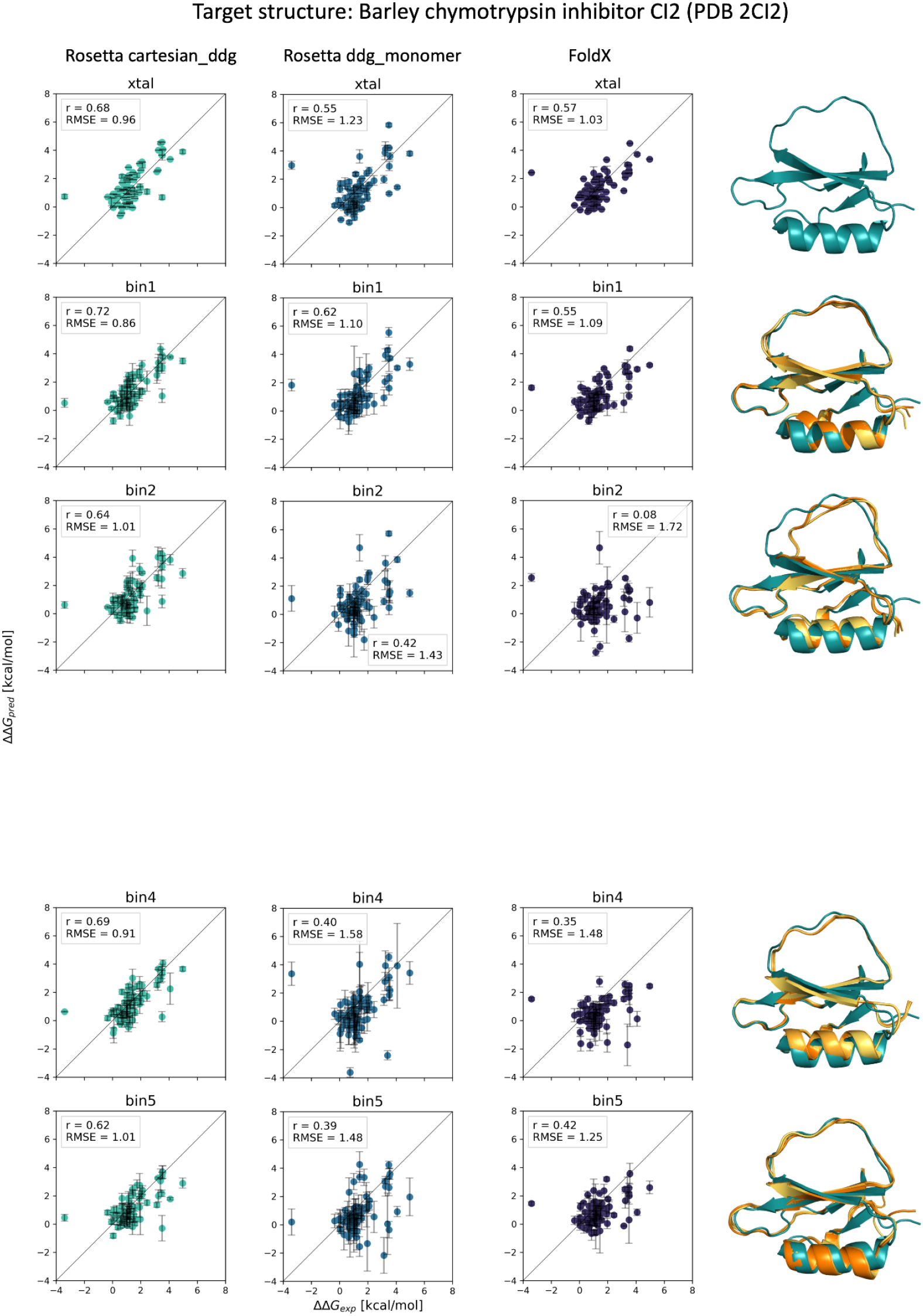
Comparison of computationally-derived ΔΔG values with experimental values on the barley chymotrypsin inhibitor CI2 using three different ΔΔG-prediction methods and either the crystal structure or a homology model. Analogous to Figure 1 in the main manuscript. No homologs were found in bin3 (below 70% but at least 55% sequence identity).

**Supplemental Figure S3:**
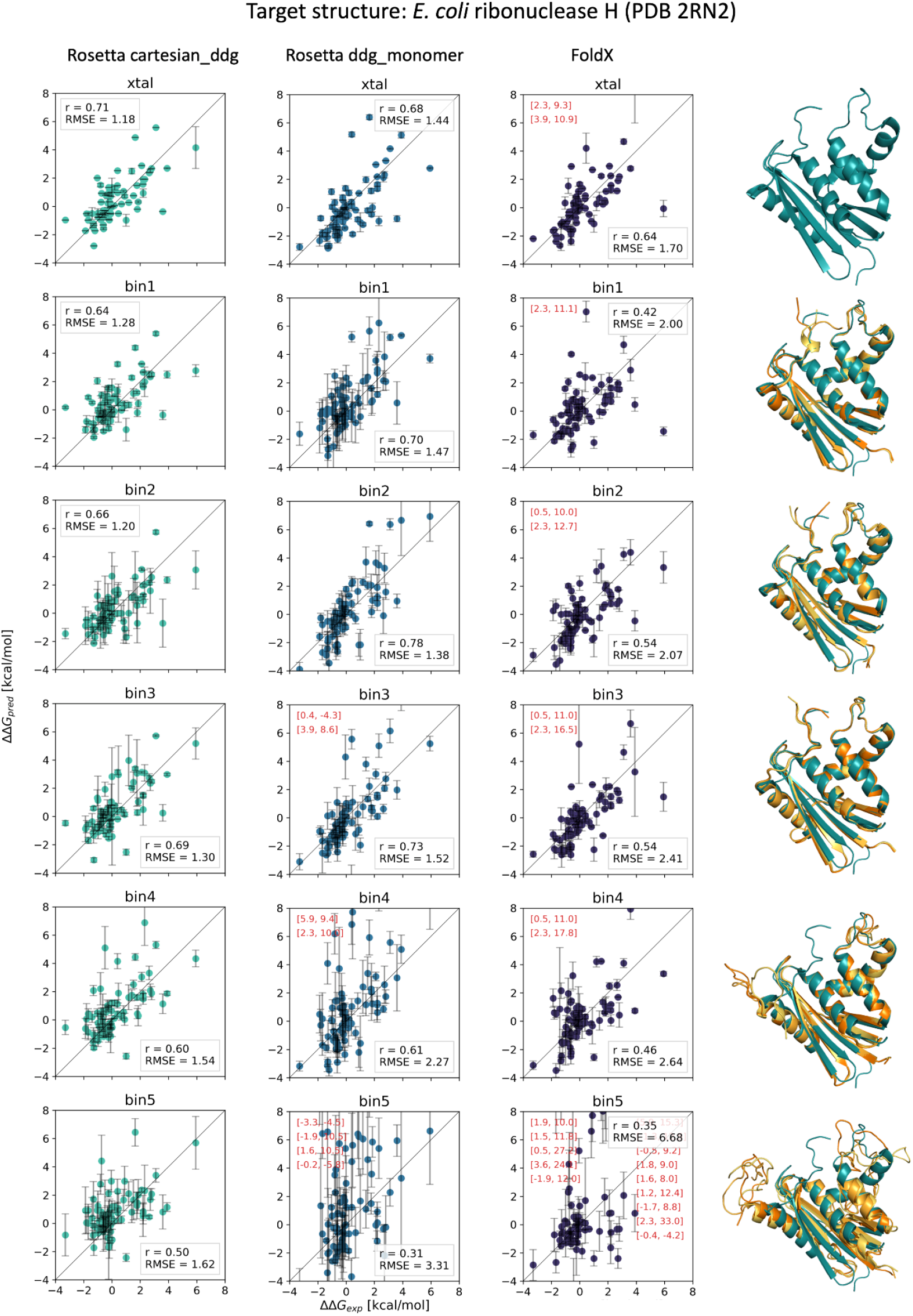
Comparison of computationally-derived ΔΔG values with experimental values on *E. coli* ribonuclease H using three different ΔΔG-prediction methods and either the crystal structure or a homology model. Analogous to Figure 1 in the main manuscript.

**Supplemental Figure S4:**
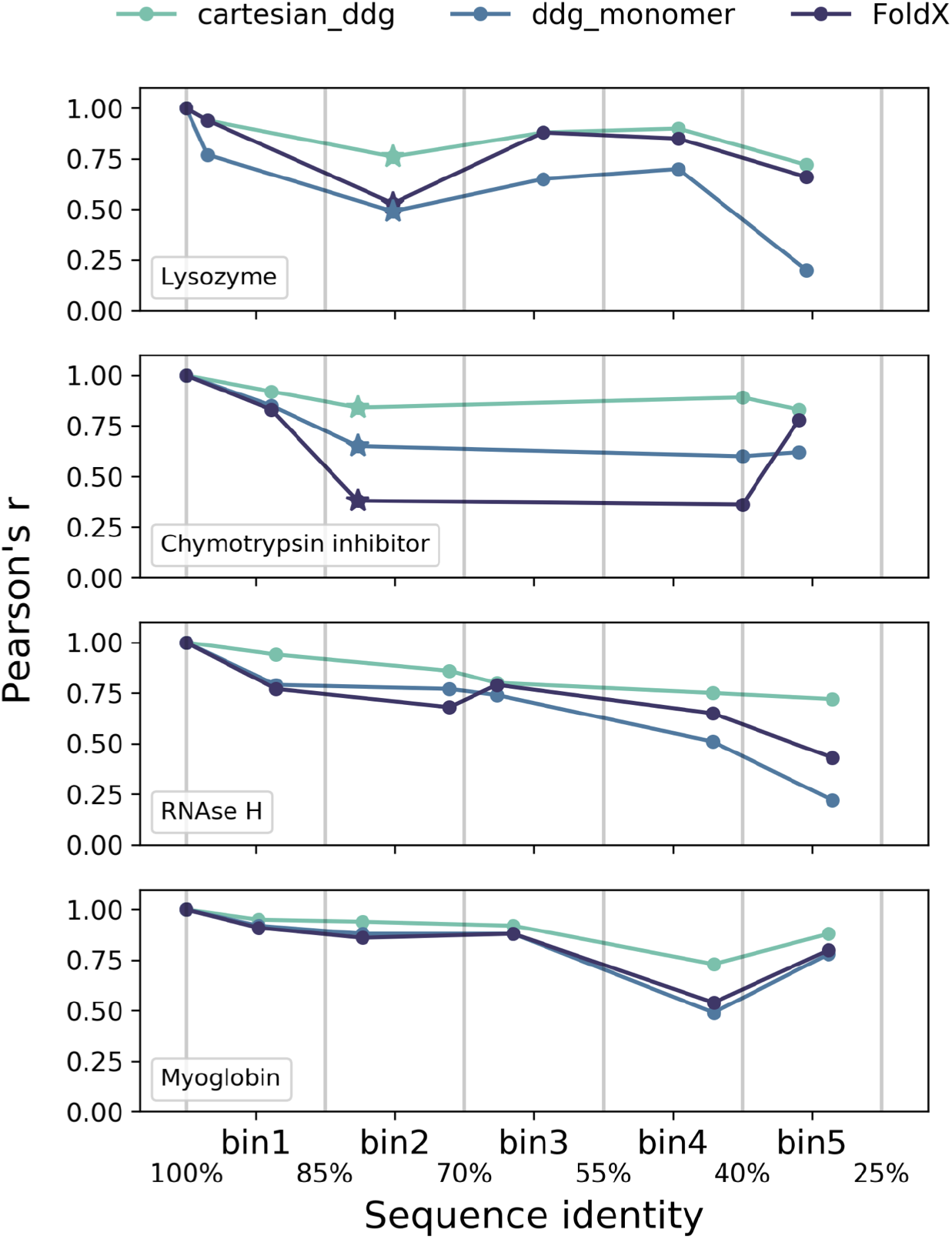
Correlations between ΔΔGs calculated on the crystal structure of the protein of interest vs. the different homology models created in this work. Pearson’s correlation coefficient. Stars in the lysozyme and chymotrypsin inhibitor plots indicate that NMR-based structures were used as templates for homology modeling in bin2.

## References

Akdel, Mehmet, Douglas E. V. Pires, Eduard Porta Pardo, Jürgen Jänes, Arthur O. Zalevsky, Bálint Mészáros, Patrick Bryant, et al. 2021. “A Structural Biology Community Assessment of AlphaFold 2 Applications.” bioRxiv. https://doi.org/10.1101/2021.09.26.461876.

Ashenberg, Orr, L. Ian Gong, and Jesse D. Bloom. 2013. “Mutational Effects on Stability Are Largely Conserved during Protein Evolution.” Proceedings of the National Academy of Sciences of the United States of America 110 (52): 21071–76.

Bienert, Stefan, Andrew Waterhouse, Tjaart A. P. de Beer, Gerardo Tauriello, Gabriel Studer, Lorenza Bordoli, and Torsten Schwede. 2017. “The SWISS-MODEL Repository-New Features and Functionality.” Nucleic Acids Research 45 (D1): D313–19.

Chung, S. Y., and S. Subbiah. 1996. “A Structural Explanation for the Twilight Zone of Protein Sequence Homology.” Structure 4 (10): 1123–27.

Dill, K. A. 1990. “Dominant Forces in Protein Folding.” Biochemistry 29 (31): 7133–55.

Fowler, Douglas M., Jason J. Stephany, and Stanley Fields. 2014. “Measuring the Activity of Protein Variants on a Large Scale Using Deep Mutational Scanning.” Nature Protocols 9 (9): 2267–84.

Frenz, Brandon, Steven M. Lewis, Indigo King, Frank DiMaio, Hahnbeom Park, and Yifan Song. 2020. “Prediction of Protein Mutational Free Energy: Benchmark and Sampling Improvements Increase Classification Accuracy.” Frontiers in Bioengineering and Biotechnology 8 (October): 558247.

Gerasimavicius, Lukas, Xin Liu, and Joseph A. Marsh. 2020. “Identification of Pathogenic Missense Mutations Using Protein Stability Predictors.” Scientific Reports 10 (1): 15387.

Goldenzweig, Adi, and Sarel J. Fleishman. 2018. “Principles of Protein Stability and Their Application in Computational Design.” Annual Review of Biochemistry 87 (June): 105–29.

Gromiha, M. M., J. An, H. Kono, M. Oobatake, H. Uedaira, and A. Sarai. 1999. “ProTherm: Thermodynamic Database for Proteins and Mutants.” Nucleic Acids Research 27 (1): 286–88.

Guerois, Raphael, Jens Erik Nielsen, and Luis Serrano. 2002. “Predicting Changes in the Stability of Proteins and Protein Complexes: A Study of More than 1000 Mutations.” Journal of Molecular Biology 320 (2): 369–87.

Johansson, Kristoffer E., Nicolai Tidemand Johansen, Signe Christensen, Scott Horowitz, James C. A. Bardwell, Johan G. Olsen, Martin Willemoës, et al. 2016. “Computational Redesign of Thioredoxin Is Hypersensitive toward Minor Conformational Changes in the Backbone Template.” Journal of Molecular Biology 428 (21): 4361–77.

Jumper, John, Richard Evans, Alexander Pritzel, Tim Green, Michael Figurnov, Olaf Ronneberger, Kathryn Tunyasuvunakool, et al. 2021. “Highly Accurate Protein Structure Prediction with AlphaFold.” Nature 596 (7873): 583–89.

Kellogg, Elizabeth H., Andrew Leaver-Fay, and David Baker. 2011. “Role of Conformational Sampling in Computing Mutation-Induced Changes in Protein Structure and Stability.” Proteins 79 (3): 830–38.

Kryshtafovych, Andriy, Alessandro Barbato, Bohdan Monastyrskyy, Krzysztof Fidelis, Torsten Schwede, and Anna Tramontano. 2016. “Methods of Model Accuracy Estimation Can Help Selecting the Best Models from Decoy Sets: Assessment of Model Accuracy Estimations in CASP11.” Proteins 84 Suppl 1 (September): 349–69.

Loladze, Vakhtang V., and George I. Makhatadze. 2005. “Both Helical Propensity and Side-Chain Hydrophobicity at a Partially Exposed Site in Alpha-Helix Contribute to the Thermodynamic Stability of Ubiquitin.” Proteins 58 (1): 1–6.

Marks, Debora S., Thomas A. Hopf, and Chris Sander. 2012. “Protein Structure Prediction from Sequence Variation.” Nature Biotechnology 30 (11): 1072–80.

Martí-Renom, M. A., A. C. Stuart, A. Fiser, R. Sánchez, F. Melo, and A. Sali. 2000. “Comparative Protein Structure Modeling of Genes and Genomes.” Annual Review of Biophysics and Biomolecular Structure 29: 291–325.

Matreyek, Kenneth A., Lea M. Starita, Jason J. Stephany, Beth Martin, Melissa A. Chiasson, Vanessa E. Gray, Martin Kircher, et al. 2018. “Multiplex Assessment of Protein Variant Abundance by Massively Parallel Sequencing.” Nature Genetics 50 (6): 874–82.

Nikam, Rahul, A. Kulandaisamy, K. Harini, Divya Sharma, and M. Michael Gromiha. 2021. “ProThermDB: Thermodynamic Database for Proteins and Mutants Revisited after 15 Years.” Nucleic Acids Research 49 (D1): D420–24.

Ó Conchúir, Shane, Kyle A. Barlow, Roland A. Pache, Noah Ollikainen, Kale Kundert, Matthew J. O’Meara, Colin A. Smith, and Tanja Kortemme. 2015. “A Web Resource for Standardized Benchmark Datasets, Metrics, and Rosetta Protocols for Macromolecular Modeling and Design.” PloS One 10 (9): e0130433.

O’Meara, Matthew J., Andrew Leaver-Fay, Michael D. Tyka, Amelie Stein, Kevin Houlihan, Frank DiMaio, Philip Bradley, et al. 2015. “Combined Covalent-Electrostatic Model of Hydrogen Bonding Improves Structure Prediction with Rosetta.” Journal of Chemical Theory and Computation 11 (2): 609–22.

Park, Hahnbeom, Philip Bradley, Per Greisen Jr, Yuan Liu, Vikram Khipple Mulligan, David E. Kim, David Baker, and Frank DiMaio. 2016. “Simultaneous Optimization of Biomolecular Energy Functions on Features from Small Molecules and Macromolecules.” Journal of Chemical Theory and Computation 12 (12): 6201–12.

Radusky, Leandro G., and Luis Serrano. 2021. “pyFoldX: Enabling Biomolecular Analysis and Engineering along Structural Ensembles.” bioRxiv. https://doi.org/10.1101/2021.08.16.456210.

Shen, Min-Yi, and Andrej Sali. 2006. “Statistical Potential for Assessment and Prediction of Protein Structures.” Protein Science: A Publication of the Protein Society 15 (11): 2507–24.

Shoichet, B. K., W. A. Baase, R. Kuroki, and B. W. Matthews. 1995. “A Relationship between Protein Stability and Protein Function.” Proceedings of the National Academy of Sciences of the United States of America 92 (2): 452–56.

Smith, C. K., J. M. Withka, and L. Regan. 1994. “A Thermodynamic Scale for the Beta-Sheet Forming Tendencies of the Amino Acids.” Biochemistry 33 (18): 5510–17.

Stein, Amelie, Douglas M. Fowler, Rasmus Hartmann-Petersen, and Kresten Lindorff-Larsen. 2019. “Biophysical and Mechanistic Models for Disease-Causing Protein Variants.” Trends in Biochemical Sciences 44 (7): 575–88.

The UniProt Consortium, and The UniProt Consortium. 2017. “UniProt: The Universal Protein Knowledgebase.” Nucleic Acids Research. https://doi.org/10.1093/nar/gkw1099.

Thiltgen, Grant, and Richard A. Goldstein. 2012. “Assessing Predictors of Changes in Protein Stability upon Mutation Using Self-Consistency.” PloS One 7 (10): e46084.

Tokuriki, Nobuhiko, and Dan S. Tawfik. 2009. “Stability Effects of Mutations and Protein Evolvability.” Current Opinion in Structural Biology 19 (5): 596–604.

Webb, Benjamin, and Andrej Sali. 2016. “Comparative Protein Structure Modeling Using MODELLER.” Current Protocols in Bioinformatics 54 (June): 5.6.1–5.6.37.

Yue, Peng, Zhaolong Li, and John Moult. 2005. “Loss of Protein Structure Stability as a Major Causative Factor in Monogenic Disease.” Journal of Molecular Biology 353 (2): 459–73.

